# Phenotyping replication is a major determinant of genomic predictive ability in sweet sorghum (Sorghum bicolor [L.] Moench)

**DOI:** 10.64898/2026.06.15.731123

**Authors:** Jean Rigaud Charles, Brian Rice, Geoffrey Morris, Thierry Tovignan, Gael Pressoir

**Affiliations:** Chibas, Centre Haïtien d’Innovation en Biotechnologies et pour une Agriculture Soutenable; Université Quisqueya, Port-au-Prince, Haïti; Soil and Crop Sciences, College of Agricultural Sciences, Colorado State University; University of Nebraska, Department of Agronomy and Horticulture; AgroUniQ Lab, Université Quisqueya, Port-au-Prince, Haïti

## Abstract

Genomic selection can increase the rate of genetic gain in crop breeding programs, but its effectiveness depends on the reliability of phenotypic data, the size and composition of the training population (TP), and the statistical model used to estimate genomic breeding values. These design choices are especially important in resource-limited breeding programs, where additional replication, larger TPs, and more extensive genotyping compete for the same resources. Using empirical data from a sweet sorghum [Sorghum bicolor (L.) Moench] breeding population, developed by CHIBAS, we evaluated the effects of phenotyping replication, TP size, training-validation genomic relatedness, and genomic prediction (GP) model on predictive ability (PA). Grain yield, plant height, stem weight, and total soluble solids were evaluated across three field environments. Few studies in sorghum have examined these factors together with comparable empirical rigor.

Increasing replication improved genomic heritability and PA for all traits and environments, with the largest gains observed for grain yield. Larger TPs and increased training-validation genomic relatedness also improved PA, but their effects were most significant when phenotype estimates were based on multiple replicates. Different GP models had comparable PA across all evaluated traits, with a few exceptions. These findings provide practical guidance for optimizing genomic selection in resource-limited sorghum breeding programs.

**ARTICLE SUMMARY:** Genomic selection can accelerate breeding only when the phenotypes used to train predictive models are reliable. Using a sweet sorghum breeding population evaluated in three Haitian field environments, we quantified how replication number, training population size, training-validation genomic relatedness, and prediction model affected genomic predictive ability for grain yield, plant height, stem weight, and total soluble solids. Replication increased genomic heritability and predictive ability for all traits, with the strongest effects for grain yield. Larger and more connected training populations improved prediction, particularly when evaluated with greater replication. These results provide practical guidance for resource-limited breeding programs.

**Core ideas:**

- In this empirical sweet sorghum breeding population, phenotyping replication was the dominant factor explaining variation in genomic predictive ability across traits and environments.
- The benefit of larger training populations and greater training-validation genomic relatedness increased when phenotype estimates were based on more replicates.
- Grain yield, the most environmentally sensitive trait evaluated, showed the largest response to improved replication and training-population design.
- Bayesian models, rrBLUP, and GBLUP showed similar predictive abilities across traits and environments, suggesting that phenotyping and experimental design may be more important than model complexity.

## INTRODUCTION

Genomic selection (GS) has become an important approach for accelerating genetic gain in plant breeding because it enables the prediction of genetic merit from genome-wide marker data rather than from a small number of marker-trait associations. The approach was first formalized for dense marker maps by Meuwissen et al. (2001) and has since been extended to many crop species as genotyping costs have decreased through next-generation sequencing and genotyping-by-sequencing platforms (Elshire et al., 2011; Wetterstrand, 2020; Satam et al., 2023). GS enables breeders to screen larger populations and select a smaller, more elite subset of individuals, thereby increasing selection intensity and potential genetic gain.

The realized value of GS depends on prediction ability (PA), which is influenced, in part, by marker density, genetic architecture, trait heritability, population structure, training population (TP) size, TP composition, and the quality of phenotypic data used to train prediction models (Heffner et al., 2011; Lorenz, 2013; Tayeh et al., 2015; Norman et al., 2018; Zhang et al., 2019; Alemu et al., 2024). Among these factors, three are directly controlled by breeders at the design stage: the size of the TP, the degree of genetic relatedness between training and selection candidates, and the intensity of field phenotyping. These factors are not independent. A fixed budget may restrict breeders from evaluating many genotypes with limited replication or fewer genotypes with more replicates. Therefore, the practical question is not whether more data improve prediction, but which information most improves prediction under field conditions.

TP size usually improves PA because larger populations sample more recombination events, capture more allelic diversity, and support more stable estimation of marker effects or genomic relationships (Lorenzana and Bernardo, 2009; VanRaden et al., 2009; Heffner et al., 2011; Lorenz, 2013; Technow et al., 2013; Endelman et al., 2014; Lorenz and Smith, 2015; Cericola et al., 2017; Tan et al., 2017; Lozada et al., 2019). However, increases in TP size may show diminishing returns, especially when additional individuals are weakly related to the prediction candidates or when phenotypic error remains high. Similarly, TP composition strongly affects prediction because PA tends to increase when the TP and prediction candidates are genetically related (Lorenz et al., 2011; Edwards et al., 2019; Werner et al., 2020; Fraslin et al., 2022). Designing TPs that balance diversity, relatedness, and cost is therefore an ongoing concern in applied genomic-assisted breeding.

Finally, PA is affected by the level of error in phenotype data. For highly polygenic traits strongly influenced by the environment, such as grain yield, low-precision phenotypes can limit prediction even when marker density and TP size are adequate (Falconer and Mackay, 2009). Additional replications reduce experimental error and improve the reliability of genotype means, thereby increasing the heritability of the target phenotype (Bos, 1983; Wricke and Weber, 1986; Falconer and Mackay, 2009; Lorenz, 2013; Lourenço et al., 2020; Yan, 2021). Yet replication is costly. For small breeding programs, allocating resources to replication can reduce the number of genotypes evaluated or the number of environments sampled. Empirical evidence is therefore needed to determine how replication interacts with TP size and composition under realistic breeding conditions.

Sweet sorghum [*Sorghum bicolor* (L.) Moench] is a multipurpose crop with potential value for grain, forage, stem biomass, sugar content, and bioenergy production. In Haiti, sorghum is relevant for crop diversification, food security, and the development of climate-resilient agricultural systems. Previous work has demonstrated the potential of GS for sorghum improvement in Haiti and highlighted the need to optimize GS for local breeding populations and environments (Muleta et al., 2019; Charles et al., 2024). However, empirical studies simultaneously evaluating replication intensity, TP size, TP composition, and model choice remain limited, particularly in small or resource-constrained breeding programs.

Here, we used an empirical multi-environment dataset from a CHIBAS sweet sorghum breeding population to quantify the relative contributions of various factors to genomic PA. Our objective was to identify practical design principles for improving GS in sorghum breeding programs operating under limited resources.

## MATERIALS AND METHODS

### Plant materials used in this study

The study used 250 sweet sorghum breeding lines derived from three CHIBAS sub-populations previously described by Charles et al. (2024). The first sub-population originated from recurrent phenotypic selection following a cross between CIRAD-ms3, an advanced selection from CIRAD, and ICSV 25280 from ICRISAT. This population was advanced through recurrent phenotypic selection to the S2 generation. The second sub-population consisted of multiple S0-derived families generated by crossing superior S0 plants from the CIRAD-ms3 × ICSV 25280 background to Coloudo Nevado, 00-SB-FSDT-427, IS23563, WILEY, CIR-1/OG2-4G-1G-M-M, PCR-2>723C-1-M-1, Papesek, and Dekabes (IS 23572). The third sub-population included lines derived from crosses between Papesek (Centa S3) and Dekabes (IS 23572).

### Field environments and experimental design

The 250 genotypes were evaluated in 2017 in randomized complete block designs with four blocks per location in three environments in Haiti in the West region of Haiti (18° 38’54.56“ N, 72° 17’ 59.43” W) on an Inceptisol. The three environments differed in planting date and water availability, ranging from fully irrigated to residual-moisture conditions (Table S1). Environment 1 was planted in September 2017 and irrigated weekly throughout the growing season. Environments 2 and 3 were planted in November 2017. Environment 2 was irrigated weekly; soil moisture was 77% one week after planting and did not fall below 50% during the season. Environment 3 was managed primarily under residual soil moisture, with rainfall increasing soil moisture to approximately 76% during the hard dough stage. Plots consisted of four rows, with 0.70 m between rows and 0.25 m within rows. Each plot measured 2.8 m by 3.5 m. Further details about the environments are previously published (Charles et al., 2024).

### Phenotype data collection

The population was phenotyped for four traits using all plants from the two central rows of each plot, minus plants on the ends. The following traits were collected: (i) Stem weight (kg/ha) was measured by weighing all leafless stems using a digital scale; (ii) grain yield (tons/ha) was estimated as 80% of panicle weight at physiological maturity; (iii) plant height (cm) was measured from the ground to flag leaf sheath on 10 plants selected randomly in the two central rows using a graduated polyvinyl chloride pipe marked with calibrated measurement lines; (iv) concentration of total soluble solids (°Brix) was measured with an Atago brand refractometer graduated from 0 to 33 %. A grinder was used to process all stems from each plot to extract the juice. The °Brix of the extracted juice was measured for three subsamples, and the mean value was calculated to obtain the final °Brix.

### Genotyping and marker filtering

The population was genotyped using genotyping-by-sequencing (GBS) at Kansas State University following the general approach of Elshire et al. (2011). DNA samples were digested with ApeKI and sequenced on an Illumina NextSeq500. The TASSEL-GBS pipeline version 4.3.7 was used for single nucleotide polymorphism (SNP) calling, and sequence tags were aligned to the sorghum reference genome Sbicolor_313_v3.0. A total of 160,066 raw SNPs were called. Markers were removed if they were non-biallelic, had a missing rate greater than 0.30, had a minor allele frequency less than 0.01, or had heterozygosity greater than 0.20. Missing marker genotypes in the retained dataset were imputed using marker means. After filtering and imputation, 28,785 SNPs were used for genomic analyses.

### Phenotypic adjustment and replication scenarios

In order to evaluate the effect of replication on GP accuracy, different scenarios were generated across the three environments. For each environment, each replicate was used in the following scenarios: only one replicate, six combinations of two replicates, and four combinations of three replicates. The complete dataset with four replicates was also included as the highest replication level.

Phenotypic data were adjusted for replicate (block) effects using univariate linear models fitted with the base R *lm()* function. Best linear unbiased estimates (BLUEs) of genotype performance were obtained using the following model:

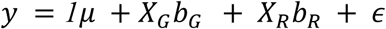

where y is the vector of BLUEs for n individuals, μ is the overall mean, *X_G_* and *X_R_* are incidence matrices for genotype and replicate effects, *b_G_* and *b_R_* are vectors of fixed effects, and *ε* is the residual term assumed to follow ε ∼ *N*(*0,Iσ^2^_ε_*). We obtained the BLUEs for the genotypic effects from the fitted model, which were used in subsequent models. For scenarios with a single replicate, genotype performance was based on a single observed phenotypic value per genotype.

### Genomic heritability estimates

Marker-based narrow-sense genomic heritability (*h^2^_g_*) was estimated for each trait, environment, and replication level using ridge regression best linear unbiased prediction (rrBLUP) implemented in the R package *rrBLUP*. Variance components were estimated from a mixed model fitted via restricted maximum likelihood (REML), with SNP effects treated as random effects. Briefly, traits were modeled as:

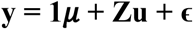

where **y** is the vector of phenotypic values for n individuals. ***μ*** is the overall mean, **Z** is an n*m design matrix containing the allele dosage at the m marker loci, **u** is an m*1 vector of m allele substitution effects, and **ɛ** is the n*1 vector of residual effects.

Unlike GP, where marker effects are used to estimate genomic best linear unbiased predictions (GBLUPs), here the objective was variance decomposition. Therefore, *h^2^_g_* was derived from the estimated variance components obtained from the rrBLUP mixed model as:

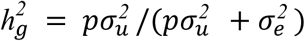

where *p* is the number of markers, *σ^2^_u_* is the marker effect variance, and *σ^2^_e_* is the residual variance. *h^2^_g_* was estimated separately for each environment and replication scenario.

### Genomic prediction and predictive ability

GP was performed using rrBLUP to estimate GBLUPs. The mixed.solve function from the rrBLUP package was used to fit the following model with a fixed effect:

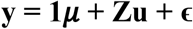

where **y** is the vector of BLUEs for n individuals. ***μ*** is the overall mean, **Z** is an n*m design matrix containing the allele dosage at the m marker loci, **u** is an m*1 vector of m allele substitution effects, and **ɛ** is the n*1 vector of residual effects. The statistical model underlying the rrBLUP approach assumes that allele substitution effects and residual effects are normally distributed (Endelman, 2011):

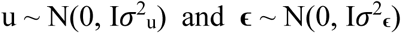

Phenotypic inputs consisted of BLUEs estimated separately for each replication scenario prior to cross-validation. Because BLUE estimation used the complete phenotypic dataset within each scenario, validation individuals contributed indirectly to variance estimation and phenotype adjustment. Consequently, the reported PAs are expected to be upwardly biased relative to fully independent prediction pipelines in which phenotype adjustment is performed separately within each training set. However, all GP scenarios evaluated in this study were affected equally by this dependency structure. Therefore, the analyses should be interpreted primarily as relative comparisons among experimental design factors (replication intensity, TP size, genomic relatedness, and prediction model) rather than as unbiased estimates of operational PA.

PA was defined as the Pearson correlation between observed phenotypic values in the validation population (VP) and predicted genomic values. Because the observed validation values are BLUEs or single-replicate observations rather than true genetic values, we use the term “predictive ability” throughout, instead of “prediction accuracy.” Since all scenarios shared the same phenotype-adjustment framework, relative differences among experimental design factors remain informative despite a possible upward bias in absolute PA. To evaluate factors affecting GP performance, three analyses were performed using the same set of phenotypic BLUEs and genome-wide SNP markers: (i) replication number, (ii) TP size, and (iii) TP composition.

### Testing the effect of phenotyping replication intensity

The effect of replication number on GP accuracy was assessed using a repeated random cross-validation approach conducted independently within each environment and scenario. For each replication level, all available genotypes were included in the analysis.

Within each environment × replication combination, genotypes were randomly partitioned into training and VPs, with 80% of individuals assigned to the training set and the remaining 20% assigned to the validation set. This random partitioning procedure was repeated 100 times to account for sampling variability and obtain robust estimates of PA. For each iteration, marker effects were estimated using the TP and subsequently used to predict genomic values for individuals in the VP. PA was calculated as the Pearson correlation between observed phenotypic BLUEs (or single-replicate observations for one-replicate scenarios) and predicted genomic values in the VP. This strategy was used to generate distributions of PA for all testing scenarios in this study.

### Testing the joint effects of TP size and phenotyping replication

The full set of 250 genotypes was randomly partitioned into five equal sized folds (50 genotypes per fold). A random seed (20241015) was set prior to assignment, and the same fold assignments were used for all traits, environments, and replication scenarios to ensure comparability.

For each iteration, one fold was randomly selected as the validation set. Four TP sizes were evaluated: 50, 100, 150, and 200 individuals, corresponding to the use of one, two, three, or four folds from the remaining four folds, respectively. Training sets were constructed by randomly sampling the required number of folds from the non-validation folds without replacement. This procedure maintains a constant validation set size (∼20% of the population) while varying training set size and ensures no overlap between training and validation. For each combination of TP size, replication level, environment, and trait, PA was evaluated over 100 independent iterations by repeating the random selection of validation fold and training folds.

### Testing the joint effects of training-validation relatedness and replication number

To evaluate the effect of TP composition, population structure was described at multiple clustering resolutions using ancestry coefficients. Sparse Nonnegative Matrix Factorization (SNMF) was used to determine the number of ancestral populations (K), using the model described by Frichot et al. (2014). The cross-entropy criterion identified K = 9 as the best-supported structure (Figure S1), and additional clustering resolutions were evaluated at K = 15, 20, 30, 40, 50, 60, 70, 80, 90, and 100. These clustering resolutions were used to generate leave-one-group-out cross-validation contrasts. For each K, each genetic group was used sequentially as the VP while all remaining groups were used as the TP. Groups with fewer than two individuals were excluded from validation. This analysis is interpreted as a test of training-validation genomic relatedness and population-structure resolution rather than as a literal modification of the underlying population. Inter-group genomic relatedness was summarized from the genomic relationship matrix by calculating the mean genomic relationship coefficient among individuals in different inferred groups at each K The genomic relationship matrix was calculated by first standardizing SNP marker genotypes and then computing pairwise Euclidean distances among individuals. These distances were transformed into genomic similarities using a Gaussian kernel function. Pairwise inter-group values were summarized to describe how genomic relatedness changed with clustering resolution. These levels of relatedness were evaluated jointly with different levels of replication number.

### Comparison of genomic prediction models

In addition to rrBLUP, five alternative GP models were compared: GBLUP, BRR, BayesA, BayesB, and BayesC. rrBLUP assumes marker effects with a common Gaussian prior variance (Endelman, 2011). GBLUP models genetic covariance through the genomic relationship matrix (VanRaden, 2008). BRR is the Bayesian analog of ridge regression implemented in BGLR (Perez and de los Campos, 2014). BayesA allows marker-specific variances, BayesB uses a mixture prior with a proportion of markers assigned zero effect, and BayesC uses a mixture prior in which non-zero marker effects share a common variance (Meuwissen et al., 2001; Habier et al., 2011). Bayesian models were run for 15,000 MCMC iterations with 3,000 burn-in iterations and thinning of 5. Model PA was estimated using repeated random 80:20 cross-validation with 100 repetitions per trait and environment, using all four replicates.

### Statistical analysis of design-factor effects

The effects of replication number, TP size, training–validation relatedness, and their interactions were evaluated using the aligned rank transform framework (Wobbrock et al., 2011), because the assumptions of residual normality and homogeneity of variance were not consistently satisfied. Analyses were performed separately by trait and environment. For each analysis, aligned-rank analysis of variance (ANOVA) was used to test main effects and interactions, and explained-variance summaries were used to compare the relative contribution of each factor. Cross-validation repetitions were used to estimate the distribution of PA for each scenario, with scenario-level summaries treated as the primary unit of inference.

## RESULTS

### Replication increased estimates of genomic heritability

Marker-based *h^2^_g_* increased with replication for all four traits in all three environments (Figure 1). Grain yield had lower *h^2^_g_* than plant height, stem weight, and total soluble solids, consistent with the higher environmental sensitivity of yield (Falconer and Mackay, 2009). For grain yield, *h^2^_g_* it increased from 0.305 with one replicate to 0.566 with four replicates in environment 1, from 0.557 to 0.891 in environment 2, and from 0.259 to 0.705 in environment 3. Total soluble solids, plant height, and stem weight generally showed higher *h^2^_g_* than grain yield, and also benefited from additional replication. These patterns indicate that replication improved the reliability of the phenotypic targets used for prediction.

**Figure 1.**
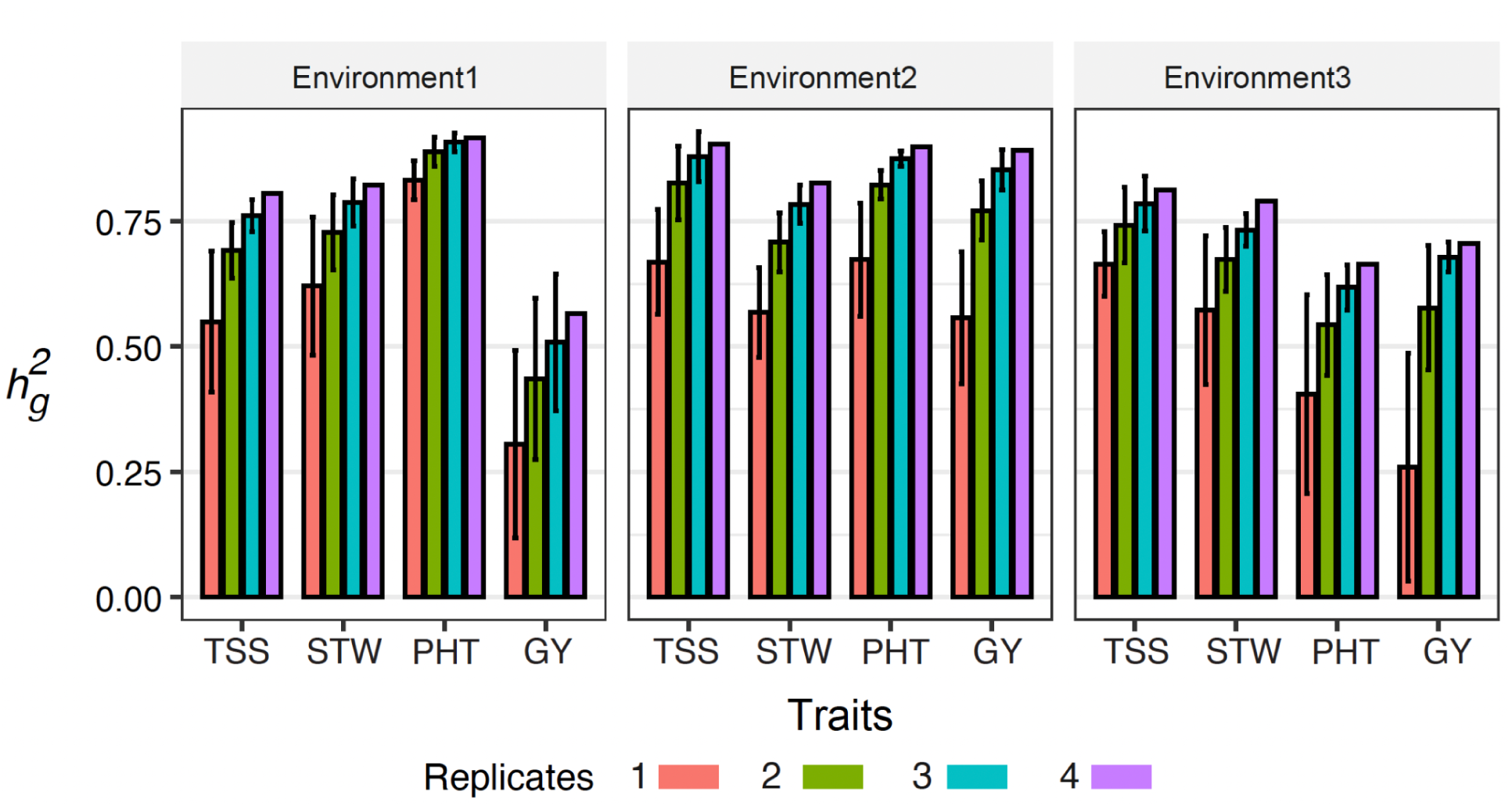
Effect of replication number on marker-based genomic heritability (*h^2^_g_*) across traits and environments. Bar height indicates the mean *h^2^_g_* estimates for grain yield (GY), plant height (PHT), stem weight (STW), and total soluble solids (TSS) in each environment, calculated using 1-4 replicates. Error bars represent the standard deviation across combinations of different observations at a given replicate level.

### Predictive ability improved with replication across traits and environments

PA increased with replication for all traits and environments (Figure 2). For total soluble solids, PA increased from 0.40 to 0.59 in environment 1, from 0.56 to 0.74 in environment 2, and from 0.52 to 0.63 in environment 3 when one and four replicates were compared. For plant height, PA increased from 0.59 to 0.71 in environment 1, from 0.40 to 0.59 in environment 2, and from 0.31 to 0.48 in environment 3. For stem weight, PA increased from 0.42 to 0.58 in environment 1, from 0.44 to 0.61 in environment 2, and from 0.37 to 0.58 in environment 3. Grain yield showed the largest relative response and increased from 0.22 to 0.41 in environment 1, from 0.27 to 0.53 in environment 2, and from 0.16 to 0.38 in environment 3.

**Figure 2.**
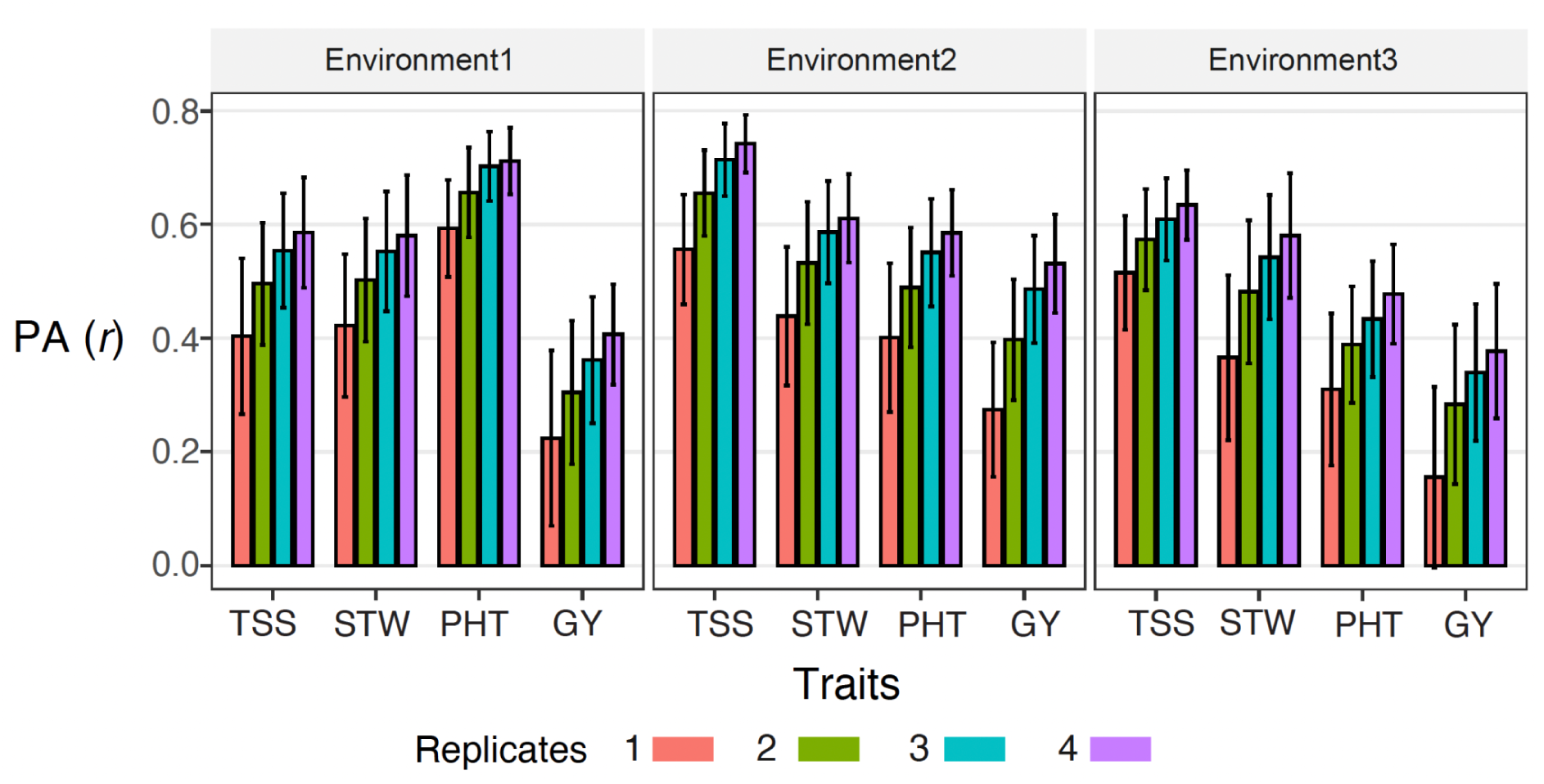
Effect of replication number on predictive ability (PA) across traits and environments. PA is the Pearson correlation (*r*) between observed genotype BLUEs or one-replicate phenotypes and predicted genomic values. Bar height indicates mean PA across cross-validation repetitions for grain yield (GY), plant height (PHT), stem weight (STW), and total soluble solids (TSS). Error bars indicate the standard deviation among the 100 cross-validation iterations.

### The benefit of increasing training population size was modest relative to increasing replication

Increasing TP size improved PA, but the benefit increased with greater replication (Figure 3). For grain yield, PA increased from 0.16 with 50 training individuals and one replicate to 0.41 with 200 training individuals and four replicates in environment 1. In environment 2, grain yield PA increased from 0.11 to 0.54 under the same contrast. In environment 3, it increased from 0.10 to 0.36 with 200 training individuals and three replicates. The aligned-rank analysis indicated that replication number was more important than TP size for grain yield PA across environments. Replication explained 88.0%, 61.7%, and 53.4% of the variation in environments 1, 2, and 3, respectively, while TP size explained 11.7%, 36.8%, and 39.1% (Table S2). Similar trends were observed in all other traits. Across traits, the response to TP size tended to plateau beyond approximately 150 individuals. Overall, the largest increases in PA were typically observed when increasing replication from one to two or three replicates, with only marginal improvements from adding a fourth replicate (Figure 3).

**Figure 3.**
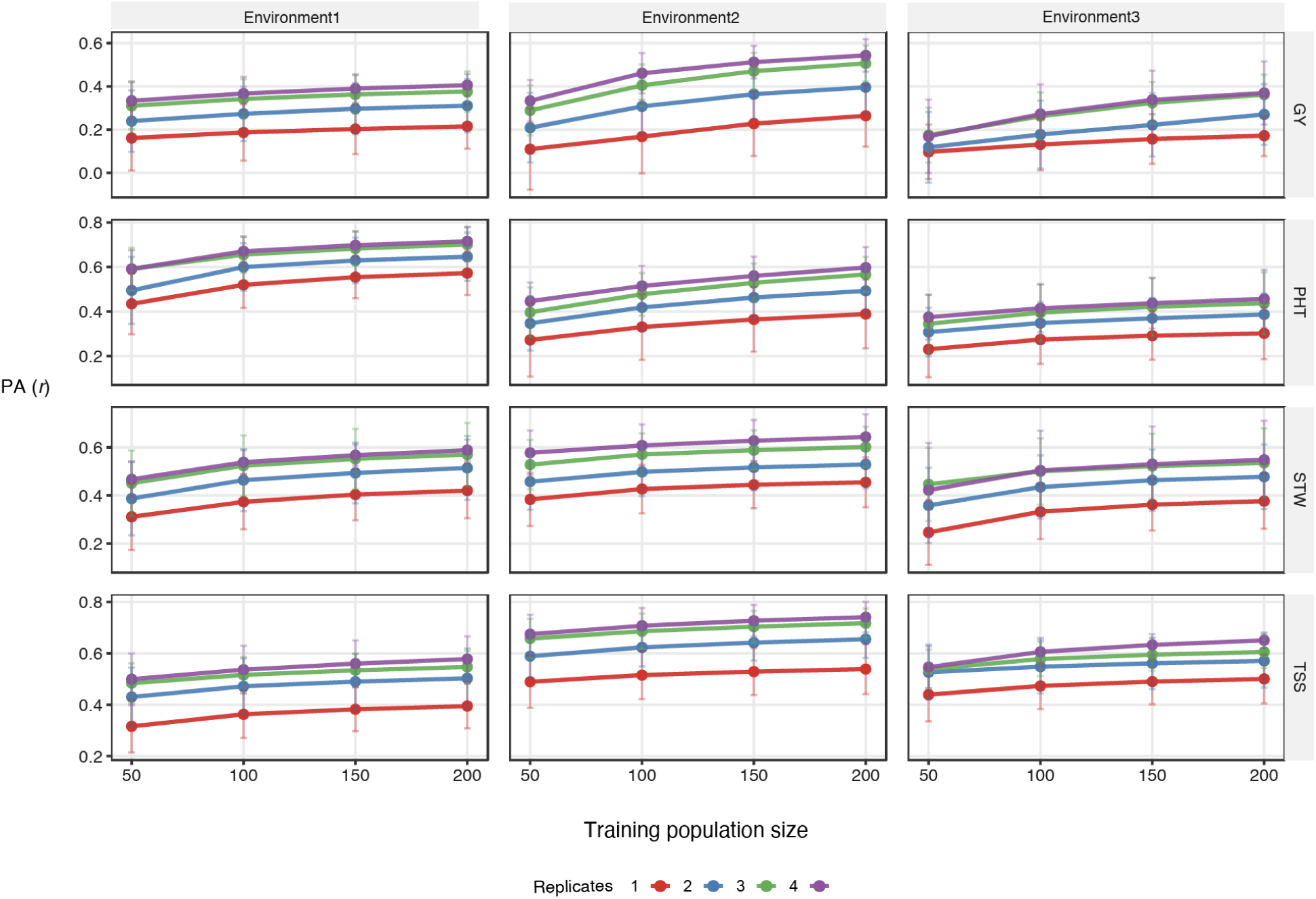
Combined effect of training population (TP) size and replication number on genomic predictive ability (PA). PA is the Pearson correlation (*r*) between observed genotype BLUEs or one-replicate phenotypes and predicted genomic values. Points represent the mean predictive ability across replications as a function of TP size for grain yield (GY), plant height (PHT), stem weight (STW), and total soluble solids (TSS). Error bars indicate the standard deviation among the 100 cross-validation iterations.

### The effect of increasing training-validation relatedness was modest compared to increasing replication

Mean inter-group genomic relatedness increased slightly as the number of inferred genetic groups increased from K = 9 to K = 100, with mean inter-group values ranging from approximately 0.35 to 0.41 (Figure S2).

PA generally increased when validation groups were more related to the TP and when phenotypes were estimated with more replicates (Figure 4). Although the increase was modest in terms of relatedness, compared to increasing the number of replicates. For grain yield, PA increased from 0.127 (K=9, 1 replicate) to 0.445 (K=90, 4 replicates) in environment 1; from 0.118 to 0.415 (K=40, 4 replicates) in environment 2; and from 0.044 to 0.238 (K=15, 4 replicates) in environment 3. Replication number explained more variation than TP composition in environments 1 (66% vs. 30%) and environment 2 (65% vs. 23%), but TP composition was more important in environment 3 (53% vs. 39%) (Table S3). For plant height, PA increased from 0.271 (K=9, 1 replicate) to 0.606 (K=70, 4 replicates) in environment 1; 0.226 (K=9, 1 replicate) to 0.515 (K=80, 4 replicates) in environment 2; and 0.208 (K=9, 1 replicate) to 0.474 (K=80, 4 replicates) in environment 3. Replication was more important in environments 2 (53% vs. 41%) and 3 (64% vs. 30%), with similar contributions in environment 1 (Table S3). For total soluble solids, PA increased from 0.153 (K=9, 1 replicate) to 0.500 (K=20, 3 replicates) in environment 1; 0.356 (K=9, 1 replicate) to 0.703 (K=60, 4 replicates) in environment 2; and 0.373 (K=9, 1 replicate) to 0.590 (K=40, 3 replicates) in environment 3 (Table S3). Replication was more important in environments 1 (58% vs. 38%) and 2 (63% vs. 34%), with similar contributions in environment 3 (Table S3). For stem weight, PA increased from 0.087 (K=9, 1 replicate) to 0.579 (K=60, 4 replicates) in environment 1; 0.264 (K=9, 1 replicate) to 0.598 (K=60, 4 replicates) in environment 2; and 0.239 (K=9, 1 replicate) to 0.526 (K=60, 4 replicates) in environment 3. Replication was more important in environments 2 (56% vs. 39%) and 3 (65% vs. 29%), with similar contributions in environment 1 (Table S3).

**Figure 4.**
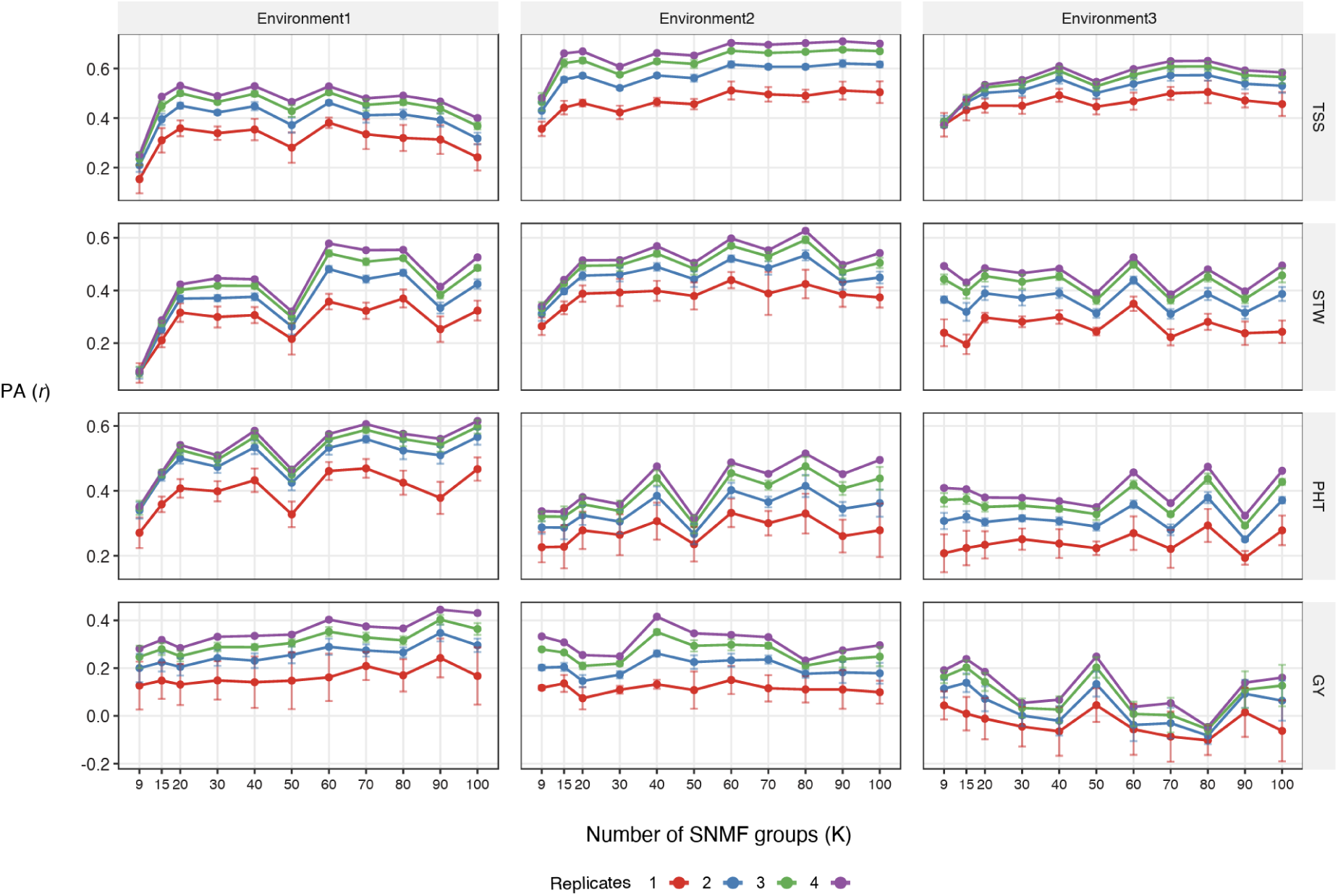
Combined effect of training-validation genomic relatedness and replication number on genomic predictive ability (PA). PA is the Pearson correlation (*r*) between observed genotype BLUEs or one-replicate phenotypes and predicted genomic values. Points show PA across sNMF clustering resolutions (K) for grain yield (GY), plant height (PHT), stem weight (STW), and total soluble solids (TSS). Higher K values represent finer population-structure partitions. Error bars indicate the standard deviation among the 100 cross-validation iterations.

### The choice of model has little impact on predictive ability

GP models produced similar PAs across traits and environments (Figure 5; Table S4). For most trait × environment combinations, there were no significant differences among models (Tukey’s HSD, *p* < 0.05). PAs generally ranged from 0.36 to 0.75, depending on the trait and environment. In Environments 2 and 3, all models showed equivalent PAs. The one expectation was that GBLUP performed significantly lower for stem weight and total soluble solids in Environment 1. Overall, these results indicate that model choice had a limited impact on genomic PA for the traits evaluated in this study.

**Figure 5.**
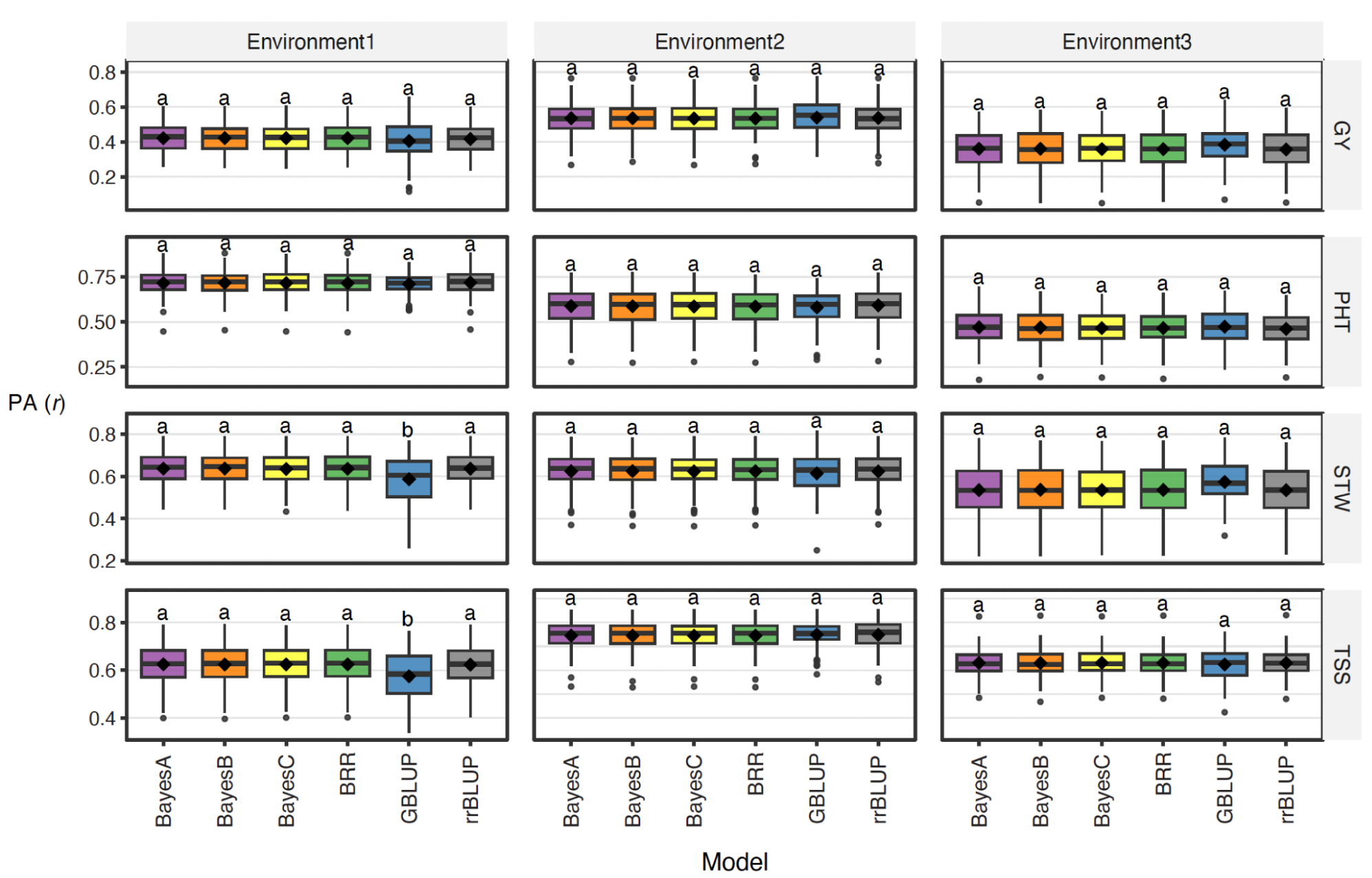
Comparison of genomic predictive ability (PA) across models. PA is the Pearson correlation (*r*) between observed genotype BLUEs and predicted genomic values. Boxplots show the distribution of PA across cross-validation iterations, and black diamonds indicate mean values across three environments for grain yield (GY), plant height (PHT), stem weight (STW), and total soluble solids (TSS). Different letters indicate significant differences among models within each trait × environment combination, as determined by ANOVA followed by HSD mean grouping (*P* < 0.05).

## DISCUSSION

This study used empirical field data from a sweet sorghum breeding population to evaluate how major design factors influence GP. The central result is that replication number was the strongest and most consistent determinant of PA across traits and environments. This finding is especially important because many GP studies emphasize marker density, model choice, TP size, and relatedness, whereas field-trial precision is often not considered. In this study, the reliability of the phenotypic estimates strongly constrained PA. It was particularly stark for grain yield and thus demonstrates how replication may have the greatest effect for traits with more measurement error (i.e., lower *h^2^_g_*).

The effect of replication was first reflected in *h^2^_g_* estimates. For all traits, *h^2^_g_* increased as more replicates were used to estimate BLUEs. This pattern is expected because replication reduces plot-level error and produces more reliable genotype means (Bos, 1983; Wricke and Weber, 1986; Lorenz, 2013; Lourenço et al., 2020; Yan, 2021). When genotype means are noisy, GP models may estimate marker effects against residual variation rather than genetic signals (Kruijer et al., 2015). Increasing replication, therefore, improves the phenotype measurement and allows marker information to be used more effectively.

TP size also improved PA, consistent with theory and empirical studies in other crops (Lorenzana and Bernardo, 2009; Heffner et al., 2011; Lorenz, 2013; Rincent et al., 2017; Norman et al., 2018; Lozada et al., 2019; Zhu et al., 2021). However, the benefit of larger TPs depended on replication. Increasing the TP from 50 to 200 individuals had the largest effect when phenotype estimates were based on multiple replicates. This result has practical implications for breeding programs with limited resources: genotyping more individuals may not produce the expected benefit if the phenotype estimates used for training are too imprecise. Conversely, increasing replication without adequate training size may also limit PA. The most efficient strategy is therefore a balanced allocation that provides enough individuals for marker-effect estimation and enough replication for reliable phenotypic targets. The precise balance will rely on the target population, trait(s), and environment.

The training-validation relatedness analysis confirmed the importance of population composition. PA increased when validation groups were more related to the training set, in agreement with prior studies (Lorenz et al., 2011; Edwards et al., 2019; Werner et al., 2020; Fraslin et al., 2022). However, this result should be interpreted with caution because relatedness was not manipulated independently. Changes in clustering resolution simultaneously altered validation-group size, group composition, and the degree of relatedness between training and validation sets. Consequently, the observed effects cannot be attributed solely to relatedness. Rather, they reflect the combined influence of relatedness and population partitioning.

The comparison among different models showed that model choice had a relatively minor effect compared with phenotyping and training design. This agrees with studies reporting that many GP models perform similarly for complex traits when marker density is high and TP size is modest (Merrick and Carter, 2021; Meher et al., 2022; Liu et al., 2025). For resource-limited breeding programs, these results suggest that investments in experimental design and phenotype quality may yield larger gains than switching among prediction models.

The study also highlights the value of empirical evaluation. Simulation studies are useful because they allow controlled manipulation of heritability, population size, genetic architecture, and relatedness (Zhong et al., 2009; Lorenz, 2013; Atefi et al., 2018). However, empirical field data include heterogeneous environments, incomplete precision, and operational constraints that define the real performance of GS in breeding programs. Results derived from simulations should be validated under field conditions, particularly when designing GS systems for small or emerging breeding programs.

A limitation of the present work is that PA was measured as the correlation between predicted genomic values and observed BLUEs or one-replicate phenotypes. This metric is appropriate for comparing scenarios, but it is not equivalent to the correlation with true genetic value. Because replication improves the precision of the validation phenotype, part of the observed gain reflects a better measurement of the target trait. This is not a weakness from a breeding perspective, because breeders select based on estimated genotype performance, but it should be recognized when interpreting the magnitude of the reported effects. A second limitation is that the economic cost of replication, genotyping, and field-plot expansion was not explicitly modeled. Future work should combine PA with cost functions to identify optimum designs for specific breeding-program budgets.

## CONCLUSION

In this empirical sweet sorghum breeding population, phenotyping replication was the most consistent driver of genomic PA across traits and environments. Additional replication increased *h^2^_g_* and improved PA, with the largest gains observed for grain yield. TP size and training-validation relatedness also improved prediction, but their benefits were greatest when phenotype estimates were based on adequate replication. By contrast, commonly used GP models differed only modestly across most traits and environments. These findings indicate that resource-limited sorghum breeding programs should prioritize field-trial precision through adequate replication before expanding TP size, optimizing genomic relatedness, or exploring more complex GP models, although all three factors contributed to PA. Optimizing these design factors can make GS more effective and more cost-efficient in practical breeding systems.

## Data availability statement

The datasets and analysis scripts generated in this study will be made available in a public repository upon publication.

## Acknowledgments

This study is made possible with the support of the American People provided to the Feed the Future Innovation Lab for Collaborative Research on Sorghum and Millet (SMIL) through the United States Agency for International Development (USAID); Program activities are partially funded by the United States Agency for International Development (USAID) under Cooperative Agreement No. AID-OAA-LA-16-00003. The contents are the sole responsibility of the authors and do not necessarily reflect the views of USAID or the United States Government. This work has also been supported by the Ministry of Agriculture, Natural Resources, and Rural Development (MARNDR) in Haiti through the Technological Innovation Program in Agriculture and Agroforestry (PITAG).

## Author contributions

J.R.C. and G.P. conceived the study and coordinated the breeding population development and field evaluations.

J.R.C. performed the phenotypic and genomic analyses.

G.M., and G.P. Funding acquisition and Resources

B.R., JR.C.., and G.P. contributed to the genomic prediction strategy, the interpretation of results, and the development of the manuscript.

B.R., G.P., and T.T. revised the draft and the final version of the manuscript.

J.R.C. wrote the initial draft of the manuscript. All authors reviewed, revised, and approved the final manuscript.

## ABBREVIATIONS

BGLR: Bayesian Generalized Linear Regression
BLUE: best linear unbiased estimate
BRR: Bayesian ridge regression
CHIBAS: Centre Haitien d’Innovation en Biotechnologies et pour une Agriculture Soutenable
GBLUP: genomic best linear unbiased prediction
GBS: genotyping-by-sequencing
GP: genomic prediction
GS: genomic selection
PA: predictive ability
rrBLUP: ridge-regression best linear unbiased prediction
SNP: single nucleotide polymorphism
TP: training population
VP: validation population.

## Supplementary figures and tables

**Figure S1.**
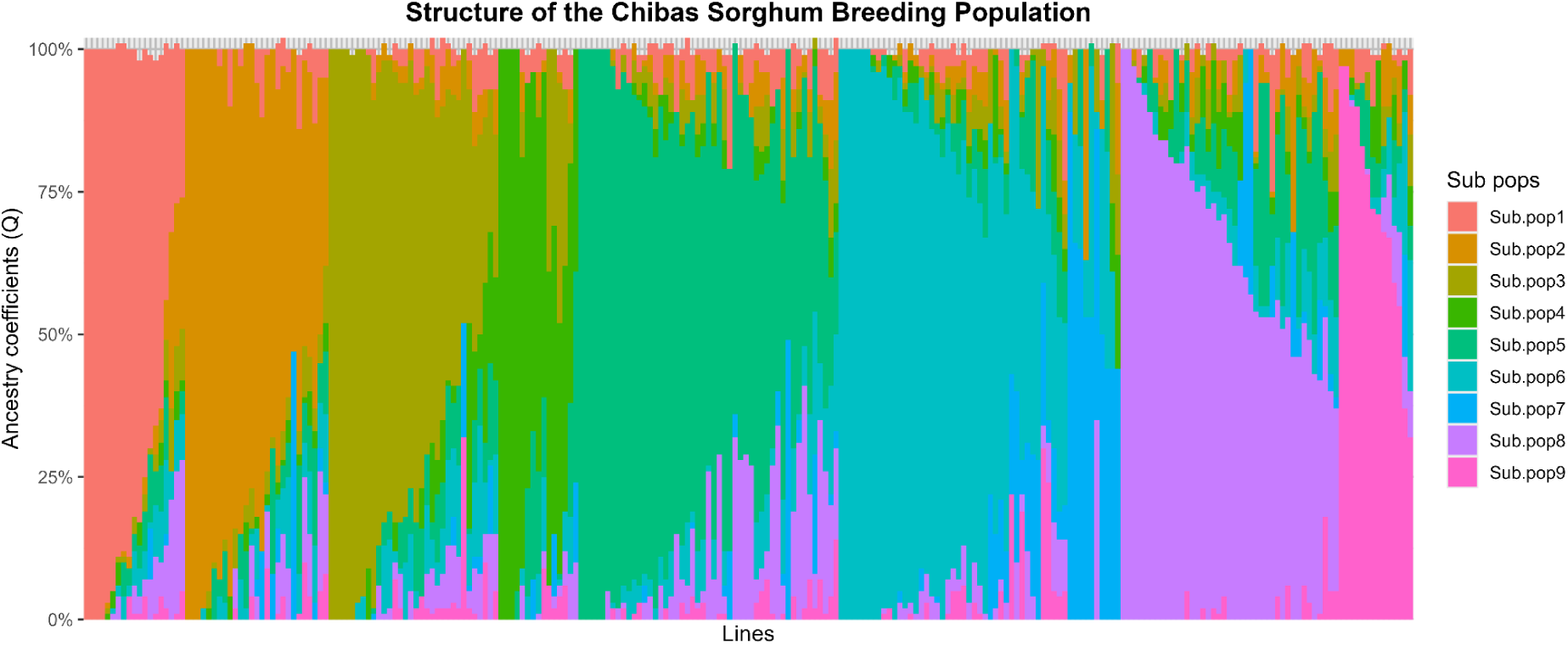
Structure of the Chibas Sorghum breeding population. The stacked bar plots show the distribution of ancestry coefficients (Q-values) for the Chibas sorghum breeding population, derived from population structure analysis. The x-axis represents individual lines of the breeding population, while the y-axis corresponds to the proportion of genetic ancestry assigned to each subpopulation (ranging from 0% to 100%). Different colors indicate the contribution of each subpopulation (Sub.pop1-9).

**Supplementary Figure S2.**
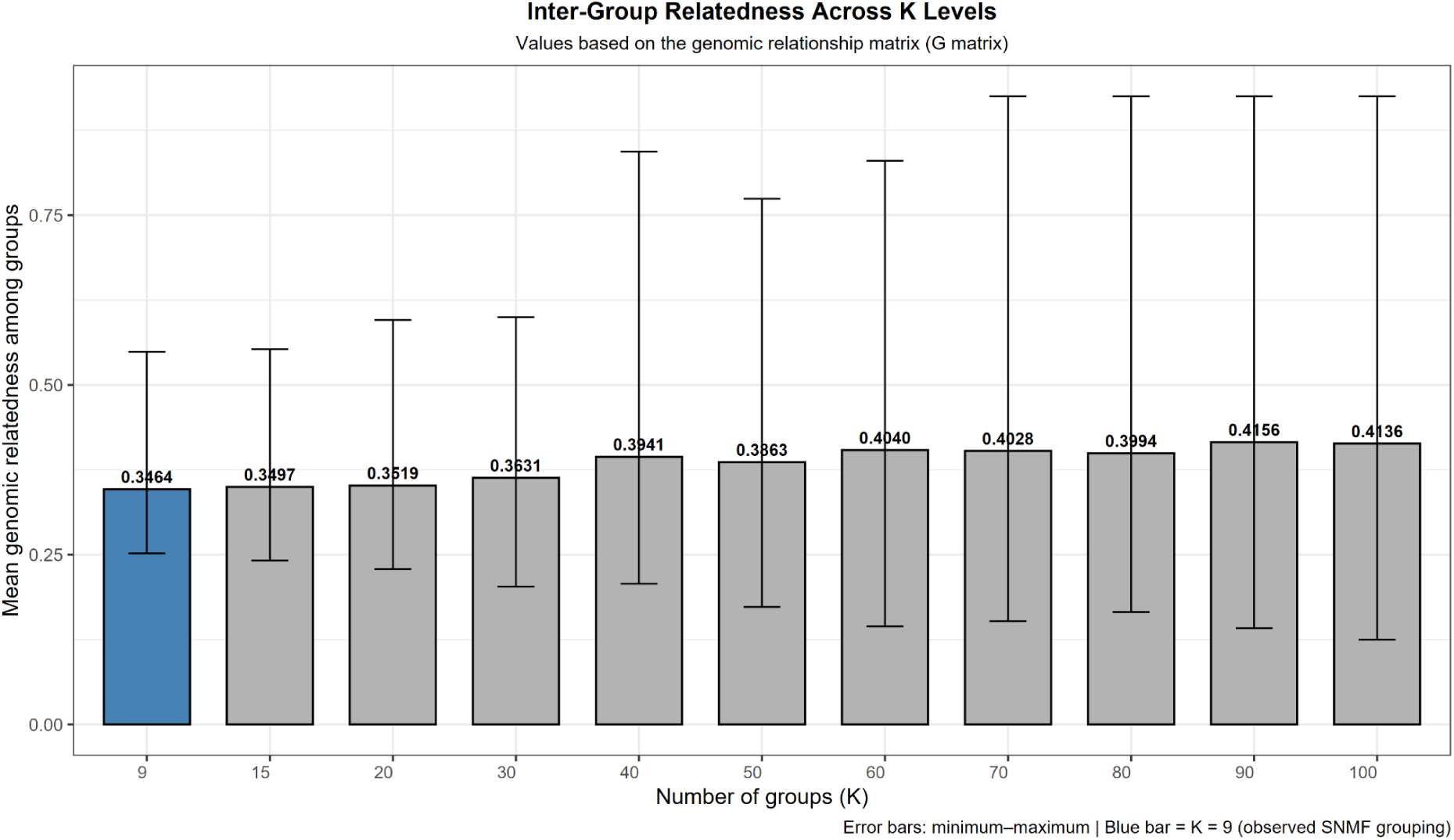
Inter-group genomic relatedness across sNMF clustering resolutions. Bar height is the mean genomic relatedness among distinct inferred groups; error bars show the minimum and maximum pairwise inter-group relatedness values.

**Supplemental Table S1.** Description of the field environments used for phenotypic evaluation.

| **Environments** | **Planting date** | **Water regime** | **Moisture context** | **Interpretive label** |
| --- | --- | --- | --- | --- |
| Environment1 | September 2017 | Weekly irrigation | Irrigated during the growing season | September irrigated |
| Environment2 | November 2017 | Weekly irrigation | High moisture; never below 50% during the season | November irrigated/high moisture |
| Environment3 | November 2017 | Residual moisture/rainfall | Rainfall increased the moisture at the hard dough stage | November residual-moisture /rainfed |

**Supplemental Table S2.** Analysis of variance for genomic prediction accuracy obtained using the ART Method (training population size and number of replications).

| **Trait** | **Environment** | **Factor** | ***P*-value** | **% Variance Explained** |
| --- | --- | --- | --- | --- |
| TSS | Environment1 | TP Size | 1.56E-06 | 16% |
| TSS | Environment1 | Replication number | 4.97E-39 | 84% |
| TSS | Environment1 | Interaction | 0.999865465 | 0% |
| STW | Environment1 | TP Size | 1.17E-14 | 34% |
| STW | Environment1 | Replication number | 3.35E-30 | 66% |
| STW | Environment1 | Interaction | 0.999989104 | 0% |
| PHT | Environment1 | TP Size | 3.64E-26 | 42% |
| PHT | Environment1 | Replication number | 6.19E-37 | 56% |
| PHT | Environment1 | Interaction | 0.887365393 | 2% |
| GY | Environment1 | TP Size | 0.002035494 | 12% |
| GY | Environment1 | Replication number | 4.30E-26 | 88% |
| GY | Environment1 | Interaction | 0.999993239 | 0% |
| TSS | Environment2 | TP Size | 3.95E-09 | 12% |
| TSS | Environment2 | Replication number | 5.72E-90 | 88% |
| TSS | Environment2 | Interaction | 0.999695912 | 0% |
| STW | Environment2 | TP Size | 1.37E-06 | 15% |
| STW | Environment2 | Replication number | 4.84E-42 | 85% |
| STW | Environment2 | Interaction | 0.999996853 | 0% |
| PHT | Environment2 | TP Size | 2.38E-19 | 42% |
| PHT | Environment2 | Replication number | 2.09E-27 | 57% |
| PHT | Environment2 | Interaction | 0.99435934 | 1% |
| GY | Environment2 | TP Size | 1.94E-31 | 37% |
| GY | Environment2 | Replication number | 1.26E-58 | 62% |
| GY | Environment2 | Interaction | 0.815553254 | 1% |
| TSS | Environment3 | TP Size | 9.06E-07 | 20% |
| TSS | Environment3 | Replication number | 1.09E-28 | 78% |
| TSS | Environment3 | Interaction | 0.904198817 | 3% |
| STW | Environment3 | TP Size | 9.16E-11 | 25% |
| STW | Environment3 | Replication number | 2.42E-34 | 74% |
| STW | Environment3 | Interaction | 0.991079807 | 1% |
| PHT | Environment3 | TP Size | 1.55E-07 | 26% |
| PHT | Environment3 | Replication number | 8.10E-23 | 74% |
| PHT | Environment3 | Interaction | 0.99939959 | 1% |
| GY | Environment3 | TP Size | 9.02E-12 | 39% |
| GY | Environment3 | Replication number | 2.94E-16 | 53% |
| GY | Environment3 | Interaction | 0.349473613 | 8% |

**Supplemental Table S3.**
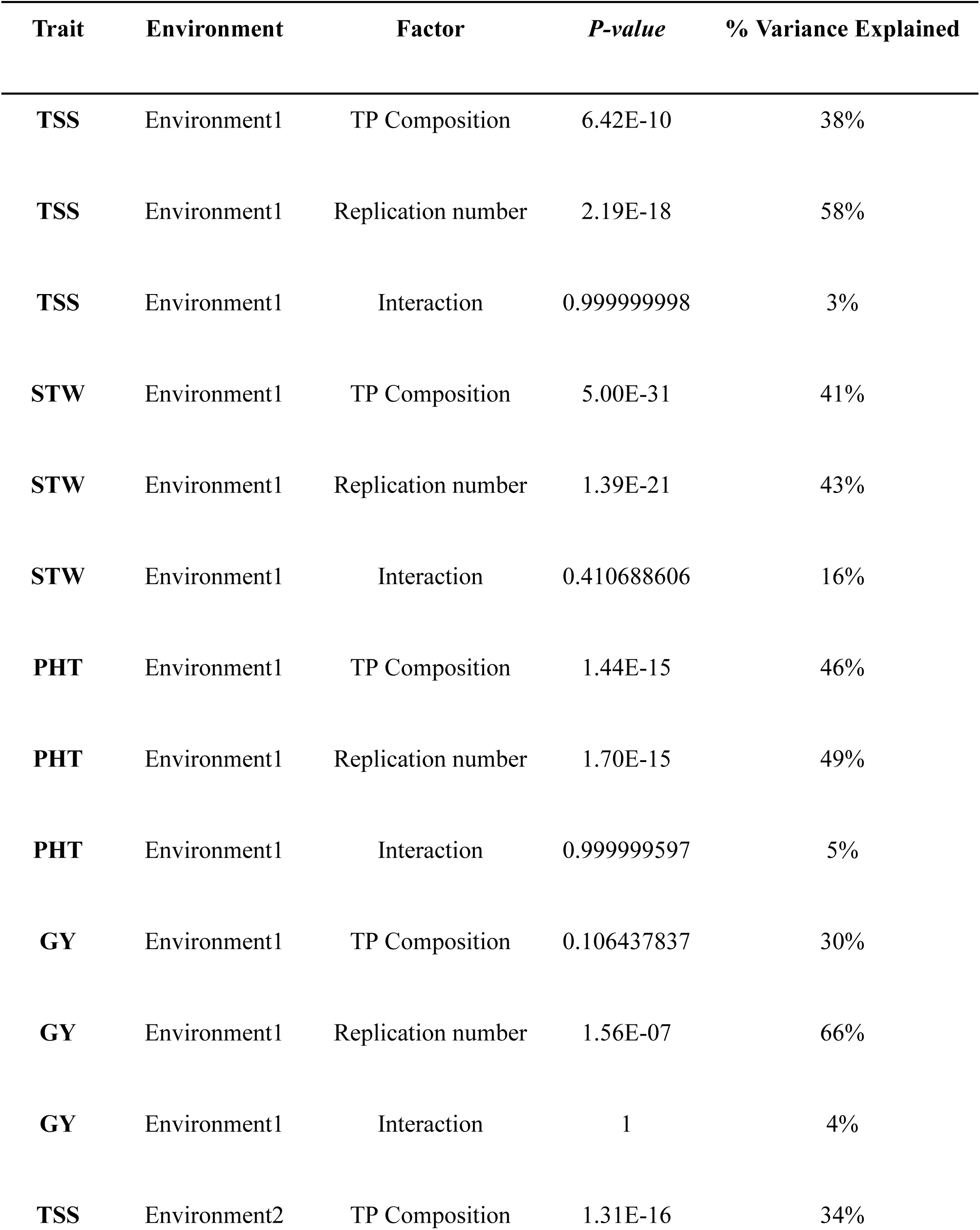
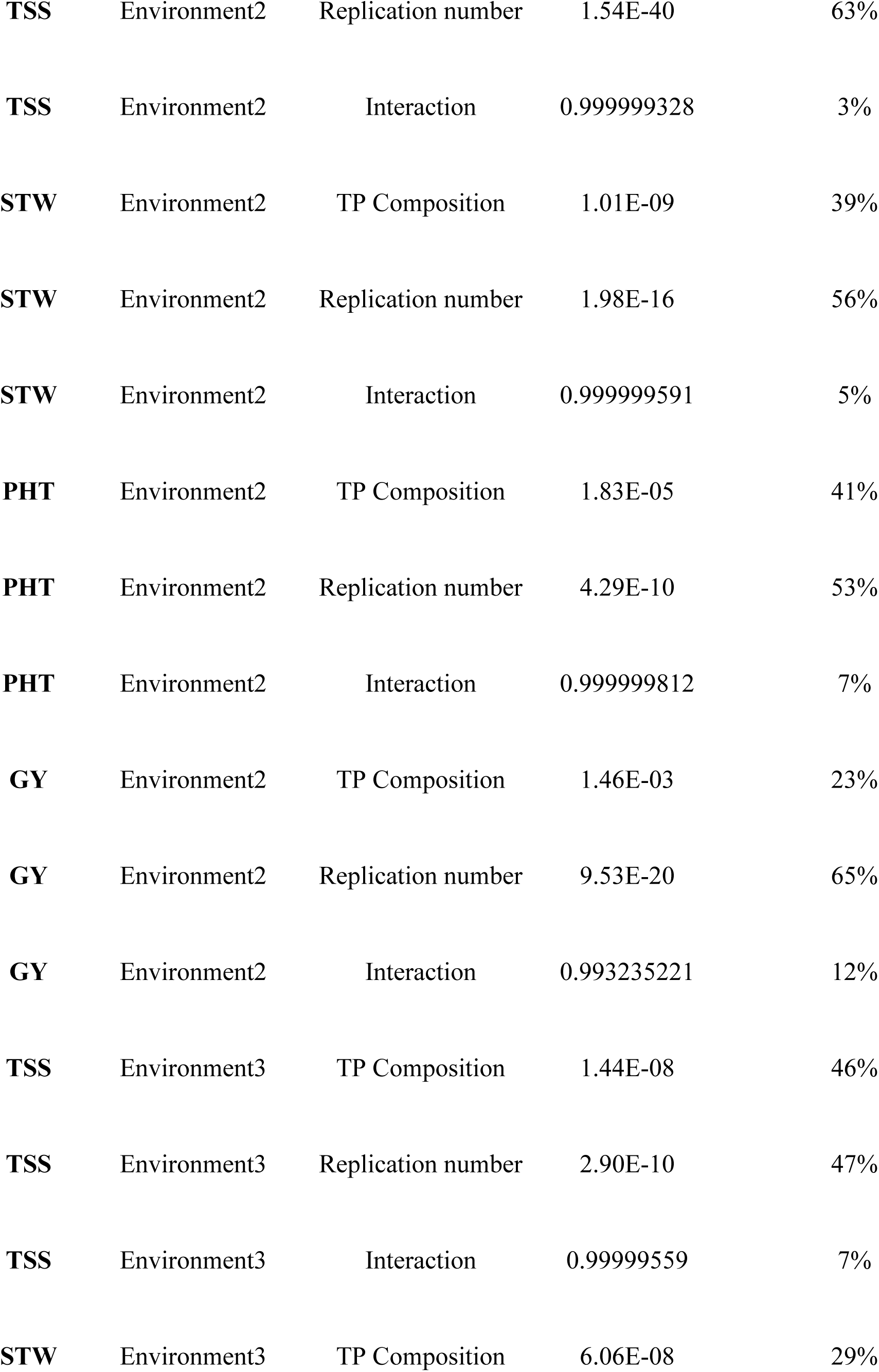
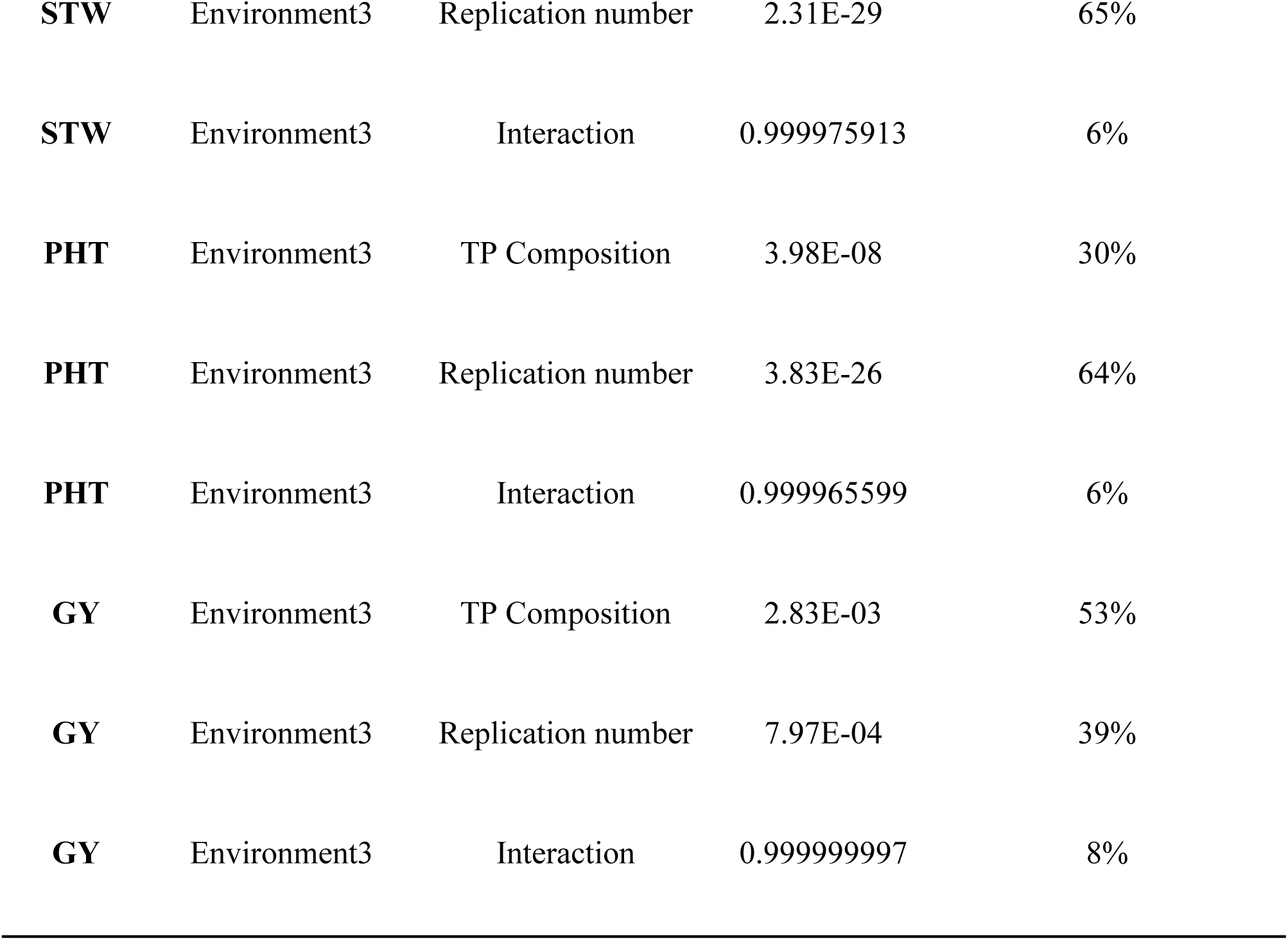
Analysis of variance for genomic prediction accuracy obtained using the ART Method (training population composition and number of replications).

| **Trait** | **Environment** | **Factor** | ***P-value*** | **% Variance Explained** |
| --- | --- | --- | --- | --- |
| **TSS** | Environment1 | TP Composition | 6.42E-10 | 38% |
| **TSS** | Environment1 | Replication number | 2.19E-18 | 58% |
| **TSS** | Environment1 | Interaction | 0.999999998 | 3% |
| **STW** | Environment1 | TP Composition | 5.00E-31 | 41% |
| **STW** | Environment1 | Replication number | 1.39E-21 | 43% |
| **STW** | Environment1 | Interaction | 0.410688606 | 16% |
| **PHT** | Environment1 | TP Composition | 1.44E-15 | 46% |
| **PHT** | Environment1 | Replication number | 1.70E-15 | 49% |
| **PHT** | Environment1 | Interaction | 0.999999597 | 5% |
| **GY** | Environment1 | TP Composition | 0.106437837 | 30% |
| **GY** | Environment1 | Replication number | 1.56E-07 | 66% |
| **GY** | Environment1 | Interaction | 1 | 4% |
| **TSS** | Environment2 | TP Composition | 1.31E-16 | 34% |
| **TSS** | Environment2 | Replication number | 1.54E-40 | 63% |
| **TSS** | Environment2 | Interaction | 0.999999328 | 3% |
| **STW** | Environment2 | TP Composition | 1.01E-09 | 39% |
| **STW** | Environment2 | Replication number | 1.98E-16 | 56% |
| **STW** | Environment2 | Interaction | 0.999999591 | 5% |
| **PHT** | Environment2 | TP Composition | 1.83E-05 | 41% |
| **PHT** | Environment2 | Replication number | 4.29E-10 | 53% |
| **PHT** | Environment2 | Interaction | 0.999999812 | 7% |
| **GY** | Environment2 | TP Composition | 1.46E-03 | 23% |
| **GY** | Environment2 | Replication number | 9.53E-20 | 65% |
| **GY** | Environment2 | Interaction | 0.993235221 | 12% |
| **TSS** | Environment3 | TP Composition | 1.44E-08 | 46% |
| **TSS** | Environment3 | Replication number | 2.90E-10 | 47% |
| **TSS** | Environment3 | Interaction | 0.99999559 | 7% |
| **STW** | Environment3 | TP Composition | 6.06E-08 | 29% |
| **STW** | Environment3 | Replication number | 2.31E-29 | 65% |
| **STW** | Environment3 | Interaction | 0.999975913 | 6% |
| **PHT** | Environment3 | TP Composition | 3.98E-08 | 30% |
| **PHT** | Environment3 | Replication number | 3.83E-26 | 64% |
| **PHT** | Environment3 | Interaction | 0.999965599 | 6% |
| **GY** | Environment3 | TP Composition | 2.83E-03 | 53% |
| **GY** | Environment3 | Replication number | 7.97E-04 | 39% |
| **GY** | Environment3 | Interaction | 0.999999997 | 8% |

**Supplemental Table S4.** Mean genomic predictive ability (*r*) of genomic prediction models across environments and traits (GY = grain yield; PHT = plant height; STW = stem weight; TSS = total soluble solids). Values followed by the same letter are not significantly different (Tukey’s HSD, *P* < 0.05).

| **Trait\*** | **Environment** | **BRR** | **BayesA** | **BayesB** | **BayesC** | **GBLUP** | **rrBLUP** |
| --- | --- | --- | --- | --- | --- | --- | --- |
| **GY** | Environment1 | 0.421a | 0.421a | 0.422a | 0.421a | 0.405a | 0.416a |
| **GY** | Environment2 | 0.534a | 0.536a | 0.536a | 0.534a | 0.539a | 0.538a |
| **GY** | Environment3 | 0.358a | 0.360a | 0.360a | 0.359a | 0.384a | 0.357a |
| **PHT** | Environment1 | 0.715a | 0.716a | 0.717a | 0.715a | 0.711a | 0.719a |
| **PHT** | Environment2 | 0.585a | 0.588a | 0.587a | 0.586a | 0.580a | 0.592a |
| **PHT** | Environment3 | 0.466a | 0.470a | 0.469a | 0.466a | 0.473a | 0.461a |
| **STW** | Environment1 | 0.636a | 0.636a | 0.637a | 0.635a | 0.586b | 0.636a |
| **STW** | Environment2 | 0.624a | 0.624a | 0.625a | 0.623a | 0.614a | 0.623a |
| **STW** | Environment3 | 0.535a | 0.535a | 0.536a | 0.535a | 0.573a | 0.534a |
| **TSS** | Environment1 | 0.625a | 0.625a | 0.624a | 0.624a | 0.575b | 0.623a |
| **TSS** | Environment2 | 0.745a | 0.745a | 0.745a | 0.744a | 0.749a | 0.748a |
| **TSS** | Environment3 | 0.631a | 0.631a | 0.630a | 0.631a | 0.624a | 0.631a |

